# Sensory neurons safeguard from mutational inheritance by controlling the CEP-1/p53-mediated DNA damage response in primordial germ cells

**DOI:** 10.1101/2022.07.19.500657

**Authors:** Simon Uszkoreit, David H. Meyer, Oded Rechavi, Björn Schumacher

**Affiliations:** Institute for Genome Stability in Ageing and Disease, Medical Faculty, University Hospital and University of Cologne, Joseph-Stelzmann-Str. 26, 50931 Cologne, Germany; Cologne Excellence Cluster for Cellular Stress Responses in Ageing-Associated Diseases (CECAD), Center for Molecular Medicine Cologne (CMMC), University of Cologne, Joseph-Stelzmann-Str. 26, 50931 Cologne, Germany; Department of Neurobiology, Wise Faculty of Life Sciences and Sagol School, Tel Aviv University, Tel Aviv 6997801, Israel

## Abstract

The genome integrity control in primordial germ cells (PGCs) is prerequisite for the inheritance of stable genomes. The PGCs in *C. elegans* are embedded in a somatic niche that regulates its DNA damage response (DDR). Here, we show that the AMPK-like kinases KIN-29 and AAK-2 are required for arresting PGCs carrying persistent DNA damage. We determined that the ASI neurons, which sense environmental conditions such as nutrient availability, secrete the TGF-beta-like ligand DAF-7 that is recognized by the DAF-1 receptor in PGCs. ASI-dependent DAF-7 signaling regulates the induction of CEP-1/p53 in the PGCs amid persistent DNA damage. Using single worm whole genome sequencing, we establish that defective ASI control of the CEP-1/p53-regulated DDR in PGCs ultimately results in the inheritance of *de novo* germline mutations. Our results indicate that sensory neurons safeguard from the inheritance of germline mutations suggesting the possibility that perception of the environment could direct genetic inheritance.

**One sentence summary:** The ASI sensory neurons regulate the CEP-1/p53-dependent DNA damage response of primordial germ cells via TGF-beta signaling and influence inherited mutational burden.

## Introduction

Genome integrity is of utmost importance for primordial germ cells (PGCs) as they generate all gametes that carry the genetic information throughout the generations. Mutations arising in PGCs can have profound effects as they are carried by all resulting germ cells. In mammals, PGCs have remained experimentally less accessible due to their complex migratory behavior through different niches during embryonic development. In contrast, in *C. elegans* L1 larvae, the germline primordium consists of only four cells in close proximity, two somatic gonad precursors (SGPs) that flank the two PGCs. PGC proliferation depends on a close communication between both cell types and the onset of proliferation is tightly regulated and demands favorable environmental conditions such as food availability (Korta and Hubbard, 2010). If conditions are unfavorable and no food is perceived, PGCs will not start to proliferate and L1 larvae instead will go into the alternative and strictly regulated dauer stage, which can last until conditions become more suitable (Golden and Riddle, 1984). This phenomenon demands strict regulation by sensors perceiving signals from environmental and nutritional cues, such as chemosensory neurons and downstream signaling pathways. Among others, the two sensory ASI neurons translate signals about food availability and pheromones into endocrine signals and influence the early dauer decision (Bargmann and Horvitz, 1991). For example, ASI neurons are responsive to Ascarosides, which are multifunctional smallmolecule signals that interfere with dauer stage and sex-specific and social behaviors. ASI neurons are the exclusive source of the DAF-7/TGF-beta ligand (Ludewig and Schroeder, 2013), which can act cell-autonomously and non-cell-autonomously to regulate expression of chemosensory receptors in the ventral nerve cord and also signals to distal tip cells of adult germlines (McGehee et al., 2015; Nolan et al., 2002; Pekar et al., 2017).

We previously showed that the PGCs remain arrested, when their genomes are damaged even though somatic development can proceed normally when the damage is confined to PGCs. As a well-defined genotoxin, UV-induced DNA damage is repaired by global-genome nucleotide excision repair (GG-NER) that is initiated by the conserved XPC-1 protein. In humans, *XPC* mutations lead to Xeroderma pigmentosum (XP) that is characterized by several thousand-fold increased skin cancer risk due to mutations arising from unrepaired UV lesions. In *C. elegans*, GG-NER is specifically required in proliferating cell types such as germ cells and in the rapidly proliferating somatic cells during early embryogenesis. In L1 larvae of *C. elegans* GG-NER is exclusively present in PGCs where it is required to remove UV-induced DNA lesions to allow the formation of the germline (Ou et al., 2019). UV irradiation of GG-NER defective *xpc-1* mutant larvae results in a permanent PGC arrest even though somatic development proceeds normally resulting in germline-less infertile adults. The PGC arrest is induced by the *C. elegans* p53-like, CEP-1, protein. We previously established that the CEP-1/p53 induction in damaged PGCs is regulated by the induction of the translation-initiation factor IFE-4 in the SGPs and determined that FGF-like signaling between the PGCs and SGPs mediates the somatic niche control of the DNA damage response (DDR) in PGCs (Figure 1A). Moreover, the IFE-4 regulated niche control of the p53 DDR is highly conserved in mammals (Ou et al., 2019).

**Figure 1:**
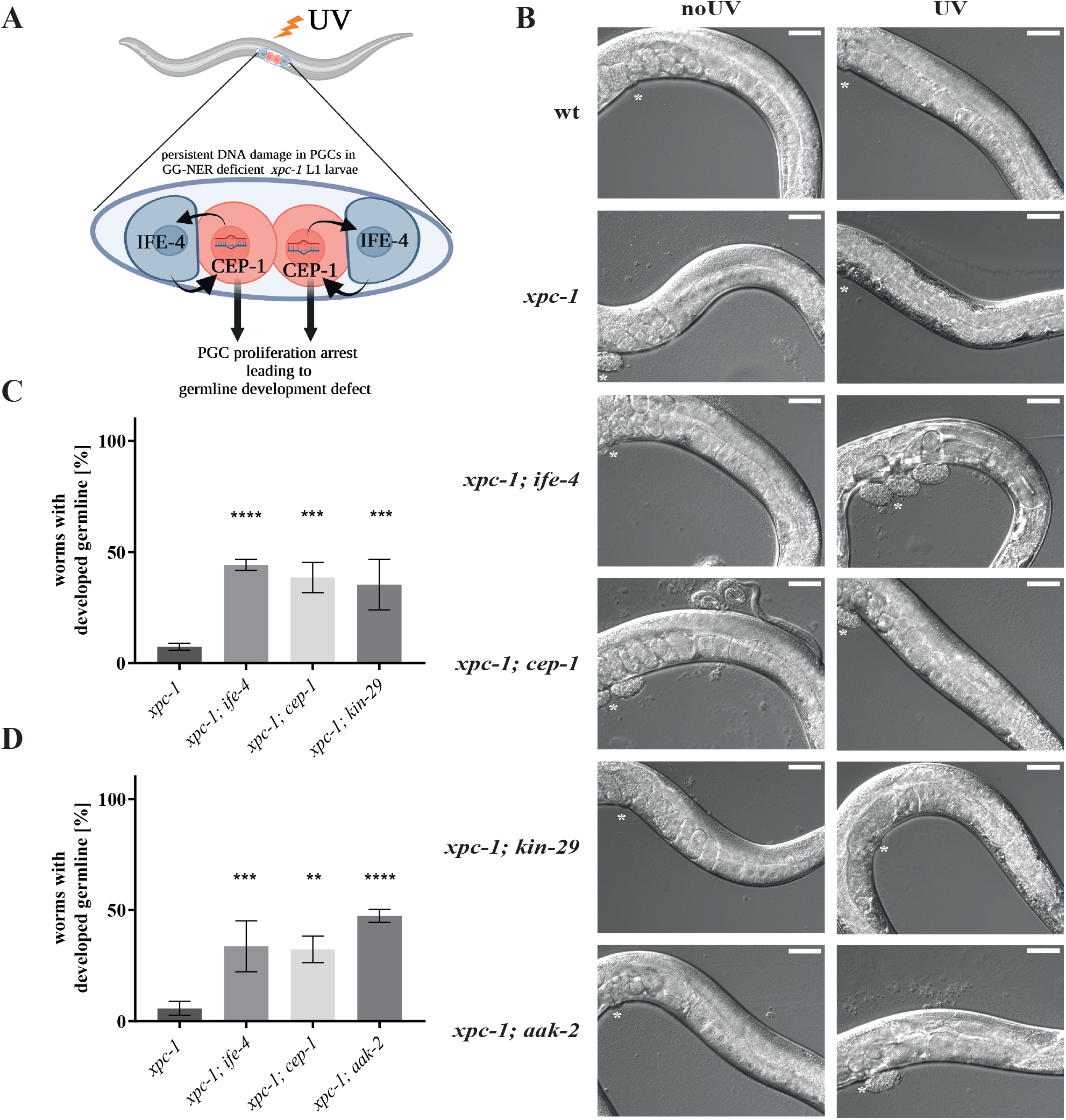
Abrogated AMPK signaling leads to restoration of germline development upon UV irradiation. **(A)** Schematic of the non-cell-autonomous DNA damage response in primordial germ cells upon persistent UV-induced damages. UV-induced DNA lesions remain unrepaired in PGCs (red) due to defective GG-NER in *xpc-1* mutant L1 larvae. The PGCs carrying persistent DNA damage communicate via FGF-like signaling to induce IFE-4 in the somatic gonad precursor (SGP) cells (blue), which is required for the CEP-1/p53 induction in PGCs that leads to DNA damage-induced cell cycle arrest resulting developed animals that lack a germline. **(B)** Representative images of germline development assay; image acquisition 3 days post mock- or UV-treatment; asterixis indicate vulva; scale bars represent 50 μm. **(C, D)** GG-NER defective AMPK mutant strains *xpc-1; kin-29* and *xpc-1; aak-2* show alleviation of germline development compared to *xpc-1*, similar to known suppressors, *xpc-1; ife-4* and *xpc-1; cep-1*. Ratio of worms with developed germlines scored 3 days post-UV irradiation. 3 independent experiments with each 3 technical replicates were summarized and shown as mean ± SD; one-way ANOVA with Dunnett’s multiple comparison test was performed; **p<0.01, ***p<0.001, ****p<0.0001.

Here, we show that the PGC arrest amid DNA damage is regulated by the ciliated amphid ASI neurons, which are interpreting nutritional and environmental states. We demonstrate that several experimental paradigms of ASI neuron inactivation result in a defective PGC arrest upon DNA damage. We identified the ASI-exclusively released transforming growth factor (TGF)-beta DAF-7 ligand that is recognized by the DAF-1 receptor in the PGCs as crucial signaling mechanism relaying the neuronal signal to the damaged PGCs. We determine that the ASI controlled TGF-beta-like signaling mediates the induction of CEP-1/p53 leading to the PGC arrest. Finally, we establish that, through regulating the DDR in the germline primordium, the ASI neurons impact the inheritance of germline mutational burden to subsequent generations. Hence, we establish a non-cell-autonomous signaling pathway that crosses the so-called Weismann barrier, that was conceptualized as a protection of the genetic information that is passed through germ cells from any somatic influences. While thus far somatic influences on germline genomes has been limited to transgenerational inherited epigenetic memories (Carpenter et al., 2021; Houri-Ze’evi et al., 2016; Moore et al., 2019; Nono et al., 2020; Posner et al., 2019), we here show that neuronal signaling could impact the inheritance of *de novo* mutations that might remain permanently in the gene pool.

## Results

### AMP-dependent kinases are involved in non-cell-autonomous DDR

In order to investigate mechanisms of PGC arrest upon DNA damage, we employed the experimental paradigm of *xpc-1* mutants that lead to persistent DNA lesions in PGCs while the somatic cells are unaffected as they use transcription-coupled (TC-) NER but do not require GG-NER to repair UV-induced lesions (Mueller et al., 2014; Ou et al., 2019). We previously determined that in response to DNA damage in PGCs, IFE-4 acts in the SGP niche to regulate the CEP-1/p53 induction in PGCs and characterized the IFE-4 target EGL-15/FGFR in the PGC-SGP signaling circuit (Figure 1A). In addition, we reported that the IFE-4 target, the Ser/Thr kinase KIN-29, contributed to the DNA damage-induced cell cycle arrest of PGCs in *xpc-1* mutated L1 larvae (Ou et al., 2019). KIN-29 bears similarity to the AMPK family of kinases and was shown to act both cell-autonomously and non-cell-autonomously in sensory regulation affecting body size and dauer decision (Lanjuin and Sengupta, 2002). To verify our previous RNA interference (RNAi) knockdown results (Ou et al., 2019), we generated the double mutant *xpc-1; kin-29*. To address the effect on the DDR in PGCs, we UV-treated synchronized *xpc-1* mutant L1 larvae and assessed the subsequent development of the germline upon completion of somatic development (Figure 1B, Representative images were taken 3 days post-treatment to compare levels of germline development). As expected, *xpc-1* mutant animals failed to develop a germline following UV treatment while a *kin-29* mutation restored germline development in a significant number of *xpc-1; kin-29* double mutants (Figure 1C; WT and single mutants in GG-NER proficient background are shown in Suppl. Fig. 1). To gauge the degree of the germline development restoration, we included the previously described double mutants *xpc-1; ife-4* and *xpc-1; cep-1*, which both significantly restored germline development post UV irradiation similarly to *xpc-1; kin-29* double mutants. In contrast to *xpc-1* single mutants, the developed germlines of WT, *xpc-1; kin-29, xpc-1; ife-4* and *xpc-1; cep-1* double mutants were all leading to oogenesis and viable progeny (Figure 1B, Suppl. Figure 1 A). The few exceptions of *xpc-1* mutants, which were nevertheless scored as ‘worms with developed germline’ (~4 – 10 %, respectively to the experiments), only presented tumorous growth or empty germline resemblances (Suppl. Figure 1E). To confirm the involvement of AMPK signaling in the non-cell-autonomous DDR, we generated an independent double mutant of the AMPK alpha subunit, AAK-2, and found a similar restoration of germline development in *xpc-1; aak-2* animals upon UV irradiation (Figure 1D, Suppl. Figure 1C). Although not looking as healthy as others, developed germlines of *xpc-1; aak-2* were leading to viable progeny. Neither GG-NER proficient WT worms nor respective single mutants showed any germline development defect upon mock or UV irradiation throughout the experiments (Suppl. Figure 1B and D). Taken together, the AMPK-like kinases KIN-29 and AAK-2 are required for the response to persistent DNA damage in PGCs.

### ASI neurons regulate PGC arrest upon persistent DNA damage

AMPKs sense cellular energy states and influence germline quiescence in the dauer stage (Fukuyama et al., 2012; Hardie et al., 1998). In *C. elegans*, sensing of energetic and nutritional states is signaled to receptive tissues through chemosensory amphid neurons. The pair of ciliated ASI neurons is involved in the interpretation of environmental cues, such as nutritional availability. Known to release exclusively one of the TGF-beta family ligands, DAF-7, ASI neurons were previously suggested to regulate dauer-stage entrance upon starvation (Ren et al., 1996) (Figure 2A). Additionally, an interplay between AAK-2 and ASI neurons was suggested to influence pro-growth and differentiation pathways (Cunningham et al., 2014). Hence, based on our finding that AMPK signaling is required for the response to persistent DNA damage in the PGCs, we set out to investigate whether ASI neurons could function as mediators in this non-cell-autonomous DDR. First, we used an ASI ablation strain, carrying a split-Caspase array under ASI-specific promoters (*oyIs84*) (Beverly et al., 2011). Proper ablation was assessed by monitoring the transgenic fluorescent GFP expression under *gcy-27* promoter in ASK, ASI and ASJ neurons until both ASI neurons were fully ablated. Since the GFP signal lasted for more than 20 hours, we were wondering whether the ASI ablation occurred already in early larval stages prior to our UV-treatment or not. To test whether in this strain the ASI neurons were functionally ablated in L1 animals by the time we applied the UV treatment, we performed lipophilic DiI-staining that is only taken up by living cells. Indeed, the ASI neurons in *oyIs84* transgenic animals were not able to take up the dye already in the first hours after L1 hatching, thus confirming their functional ablation (Suppl. Figure 2 A and B). Strikingly, the ablation of ASI neurons in *xpc-1; oyIs84* mutants led to a pronounced alleviation of germline development defect post-UV compared to *xpc-1* mutants (Figure 2B).

**Figure 2:**
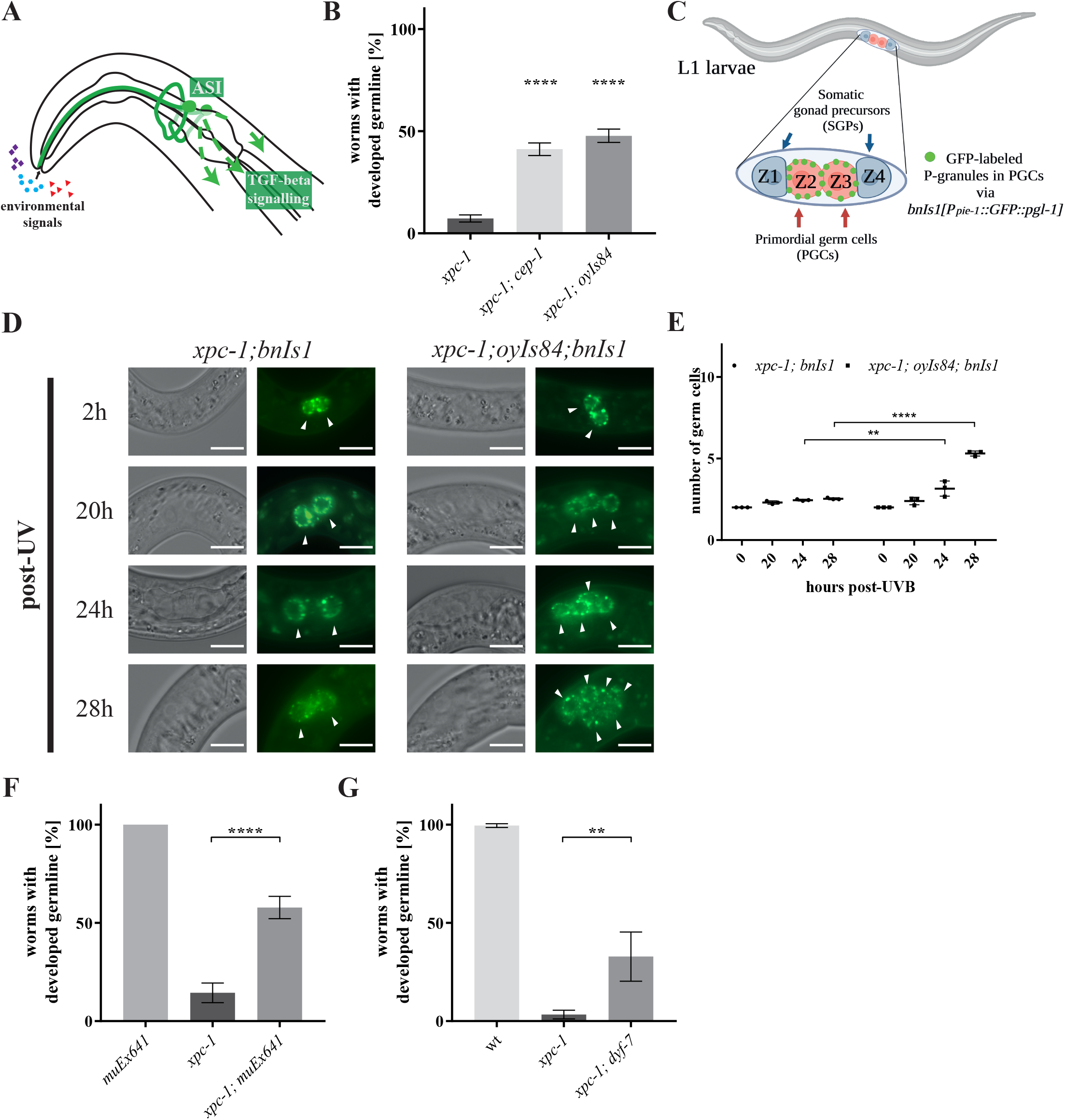
ASI neurons regulate the DNA damage response in primordial germ cells. **(A)** Schematic overview of the pair of amphid ASI neurons that respond to environmental cues by secreting the TGF-beta ligand DAF-7. **(B)** Ablation of ASI neurons via the transgene *oyIs84[gpa-4p::TU#813* + *gcy-27p::TU#814* + *gcy-27p::GFP* + *unc-122p::dsRed]*, carrying an ASI-specific split-Caspase array alleviate the impaired germline development upon UV treatment similarly to CEP-1/p53 deficiency. **(C)** Schematic overview of the primordial germline in *C. elegans* L1 larvae containing of two PGCs, Z2 and Z3, flanked by somatic gonad precursor (SGP) niche cells. PGCs can be specifically visualized with GFP-labeled P-granules via transgene *bnIs1*[P_*pie-1*_::GFP::*pgl-1* + *unc-119(+)*]. **(D)** Representative images of *xpc-1* mutant strain expressing transgene(s) *bnIs1*[P_*pie-1*_::GFP::*pgl-1* + *unc-119(+)*] or *bnIs1*[P_*pie-1*_::GFP::*pgl-1* + *unc-119(+)*]; *oyIs84[gpa-4p::TU#813* + *gcy-27p::TU#814* + *gcy-27p::GFP* + *unc-122p::dsRed]*. Arrow heads indicate PGCs; scale bars represent 10 μm. **(E)** ASI ablation alleviates PGC proliferation post UV-irradiation; combined results for PGC proliferation kinetics of 3 independent experiments are shown as dot blot with mean ± SD; RM two-way ANOVA with matched values followed by Sidak’s multiple comparison test was performed; **p<0.01, ****p<0.0001. **(F)** Silencing of ASI neurons via Tetanus toxin light chain expression under ASI-specific promoter on transgene muEx641[P_*gpa-4*_::GFP::Tetx + *Punc-122*::RFP] in *xpc-1;muEx641* mutant alleviates germline development defect upon UV treatment. Only transgenic RFP-positive animals, indicative of transgenic animals, were scored. **(G)** Amphid-neuron development mutant, *xpc-1; dyf-7*, shows alleviated germline development defect upon UV treatment. (B, F, G) Ratio of worms with developed germlines scored 4 days post-UV irradiation and incubation at 15 °C. 3 independent experiments with each 3 technical replicates were summarized and shown as mean ± SD; one-way ANOVA with Dunnett’s multiple comparison test was performed; **p<0.01, ****p<0.0001.

We previously established that the germline development defect in UV-treated *xpc-1* mutants is a result of the permanent PGC arrest leading to the inability to generate germ cells. To investigate the effect of ASI neuron ablation to the PGC arrest amid DNA damage, we next investigated the proliferation kinetics of PGCs. To this end, we employed a strain carrying the transgene *bnIs1*[*pie-1p::GFP::pgl-1* + *unc-119*(+)], which labels P-granules with GFP exclusively in germ cells (Cheeks et al., 2004) thus allowing tracking PGCs and subsequently proliferating germ cells (Figure 2C).

First, we used this strain to confirm that PGC proliferation in ASI deficient animals was normally arrested upon starvation to preclude the possibility that the elevated PGC proliferation kinetics were a result of a failure to properly arrest prior to feeding. We monitored the PGC cell cycle arrest in worms carrying the ASI ablation transgene over time during starvation. Even after 18 hours, PGCs were not able to escape the starvation-induced cell cycle arrest (Suppl. Figure 3A) and can be only initiated after 3 to 4 hours following feeding (Fukuyama et al., 2006). Indeed, in the absence of UV irradiation germline development was indistinguishable from WT animals.

We next monitored the proliferation of PGCs following UV-induced DNA damage in *xpc-1* and *xpc-1; oyIs84* mutant animals using the *bnIs1* P-granule marker transgenic strain background. Consistent with the germline development phenotype, the DNA damage-induced proliferation arrest of the two PGCs was alleviated as, in contrast to *xpc-1* mutants, ASI ablated *xpc-1; oyIs84* mutants showed germ cell proliferation within 24 h post UV-treatment (Figure 2D and E).

To independently confirm the involvement of ASI neurons in the DNA damage-induced PGC arrest, we used a distinct paradigm of ASI-disabling, a strain carrying the transgene *muEx641* (Chisnell and Kenyon, 2018). This transgene expresses ASI-specifically a tetanus-toxin light-chain (Tetx) leading to silencing of ASI neuronal signaling by blocking the release of clear-core and dense-core vesicles but not affecting gap junctions. Since *muEx641* is extrachromosomal and can get lost throughout generations, only animals expressing the transgenic RFP co-marker were scored for germline development. Here, we observed that silenced ASI neurons led to restoration of germline development in the *xpc-1* background following UV-induced DNA damage (Figure 2F). As yet another ASI ablation strategy, we employed an overall amphid neuron developmental mutant, *dyf-7*. Importantly, this strategy provides animals deprived of ASI neurons due to a genetic mutation and independent of the usage of any transgenes. Also *dyf-7* mutants showed a restoration of germline development albeit to a lesser extent than the specific ASI ablation and silencing probably due to the drastic neuronal impairment (Figure 2G). GG-NER proficient WT animals and single mutants showed no phenotype (Suppl. Figure 3B). These results indicate that ASI neurons are required for the arrest of PGCs that carry persistent DNA damage.

The PTEN-orthologue DAF-18 has previously been implicated in germline proliferation and germline stem cell quiescence, especially upon nutritional unfavorable conditions via AMPKs and TGF-beta signaling (Fukuyama et al., 2006, 2012; Narbonne et al., 2017; Tenen and Greenwald, 2019). To investigate whether DAF-18 could be involved in the PGC DDR, we examined the germline development of *xpc-1; daf-18* mutants. However, the double mutant displayed similar germline development defects post UV as *xpc-1* single mutants ruling out an involvement of DAF-18/PTEN (Suppl. Figure 3B and C).

### TGF-beta ligand DAF-7 signals from ASI neurons to PGCs to regulate DNA damage-induced germline development arrest

We next sought to identify the ASI signal that regulates the PGC cell cycle arrest upon DNA damage. We hypothesized that the TGF-beta pathway might be involved as the TGF-beta-like ligand DAF-7 is exclusively expressed and released by ASI neurons via large dense-core vesicles to signal to downstream cells (Inoue and Thomas, 2000). Further, a connection between TGF-beta and AMPKs KIN-29 and AAK-2 has been reported (Koga et al., 1999; Zhang et al., 2019) and DAF-7 is known to signal upon unfavorable conditions to induce the dauer stage (Gallagher et al., 2013). Moreover, TGF-beta signaling via DAF-7 was shown to have the ability to act non-cell-autonomously on distal tip cells (DTCs) in adult germlines to negatively regulate expression of DAF-3 and DAF-5 (Pekar et al., 2017). To investigate the expression pattern of DAF-7 protein, we inserted a V5 Tag into the endogenous *daf-7* locus to generate a C-terminal fusion of the DAF-7 protein with the V5 Tag. Similarly to the previously established transcription report of the *daf-7* promoter driven GFP (Suppl. Fig 4A), we observed the DAF-7::V5 expression specifically in ASI neurons (Figure 3A, Suppl. Fig. 4B). Expression of DAF-7::V5 in ASI neurons was significantly increased in GG-NER deficient background, *xpc-1; daf-7::V5*, upon UV (Figure 3 A and B) indicating that DAF-7 in ASI neurons is induced in response to persistent DNA damage in PGCs.

**Figure 3:**
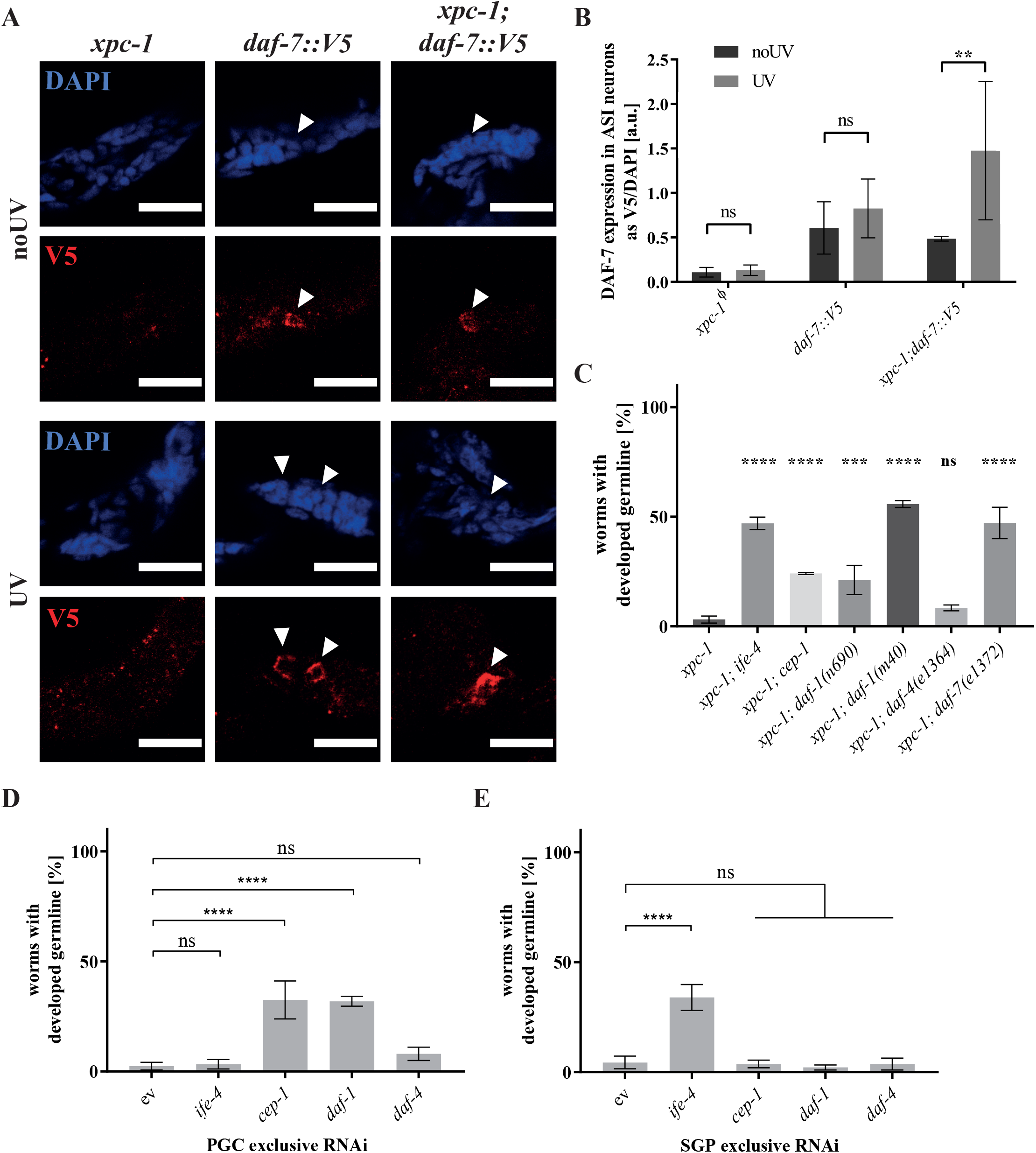
ASI-secreted TGF-β signaling mediates DNA damage response in germ cells. **(A)** Representative images of DAF-7 expression in ASI neurons via V5 staining; *xpc-1* mutant serves as staining control; arrow heads indicate ASI neuron; scale bars represent 10 μm. **(B)** Quantification of DAF-7 expression as V5/DAPI ratio in arbitrary units shows upregulation of DAF-7 post-UV treatment; Φ = cells from head region were quantified for *xpc-1* control animals; 3 independent experiments were summarized and shown as mean ± SD; two-way ANOVA with Sidak’s multiple comparison test was performed; ns = not significant, **p<0.01. **(C)** Impairment of TGF-β signaling in the *xpc-1* mutant background, in TGF-β ligand mutants *daf-7(e1372)* or TGF-β receptor 1 hypomorph *daf-1(n690)* or null *daf-1(m40)* mutants alleviate the germline development defect upon UV-induced DNA damage, while the TGF-β receptor 2 *daf-4(e1364)* mutant shows no effect. **(D, E)** Tissue-specific knockdown via RNA interference of TGFBR1 *daf-1* or TGFBR2 *daf-4*, respectively in PGCs in *xpc-1(tm3886); rrf-1(pk1417)* (D) or SGPs in *xpc-1(tm3886); rde-1(ne219); qyIs102[fos-1ap::rde-1* + *myo-2p::YFP* + *unc-119(+)]* (E), reveals importance of TGF-beta reception via DAF-1 specifically in PGCs upon persistent UV-induced DNA damage. RNAi against *cep-1/p53* and *ife-4* were included as controls for the DDR impairment in PGCs and somatic cells, respectively. (C, D, E) Ratio of worms with developed germlines scored 4 days post-UV irradiation. 3 independent experiments with each 3 technical replicates were summarized and are shown as mean ± SD; one-way ANOVA with Dunnett’s multiple comparison test was performed; ***p<0.01, ****p<0.0001, ns=not significant.

In order to address the functional involvement of DAF-7 mediated TGF beta signaling in the DDR in PGCs, we investigated the germline development in *xpc-1; daf-7* double mutants. Strikingly, *xpc-1; daf-7* double mutants showed a significant suppression of the germline development defect compared to *xpc-1* mutants following UV treatment (Figure 3C). To ascertain the specificity of the DAF-7 involvement in the DDR, we validated that *xpc-1; daf-7* double mutant PGCs underwent the normal PGC cell cycle arrest freshly hatched L1 larvae in starved conditions similarly to WT and *xpc-1* single mutants (Suppl. Figure 3D). The TGF-beta ligand DAF-7 is recognized by either of the canonical TGF-beta receptors DAF-1 and DAF-4. While *xpc-1; daf-1* double mutants led to a drastic increase in the fraction of animals developing a germline, the double mutant *xpc-1; daf-4(e1364)* did not suppress the germline development defects of UV treated *xpc-1* mutants. Moreover, a hypomorph *daf-1(n690)* point mutant led to a milder suppression of the germline development defects than a null *daf-1(m40)* deletion mutant in the *xpc-1* background (Figure 3C). The respective single mutants showed no particular sensitivity to UV treatment in a GG-NER proficient background (Suppl. Figure 3E).

To determine where the TGF-beta signaling is received, we used tissue-specific RNAi mutants in the *xpc-1* background. While *xpc-1; rrf-1* mutants are RNAi defective in somatic cell types but are RNAi sensitive in germ cells, RNAi proficiency in *xpc-1; rde-1; qyIs102[Pfos-1a::rde-1*] mutants is specifically restricted to the SGPs that form the somatic niche of the PGCs. We previously extensively confirmed the cell type specificity of those mutant strains also in the context of UV-induced DNA damage in L1 larvae (Ou et al., 2019). We determined that tissue-specific knock down of *daf-1* in PGCs restored germline development upon UV treatment similar to the levels of *cep-1* knock down, while no suppression of germline development arrest could be observed for SGP-specific knock down of *daf-1* (Figure 3D and E). We confirmed the SGP-specific RNAi proficiency by *ife-4* knockdown leading to a robust suppression of the germline development defect. These results show that the TGF-beta ligand DAF-7 mediates the ASI neuronal input to the DAF-1 TGF-beta receptor in PGCs to induce their arrest when they carry persistent DNA damage.

Furthermore, it was shown that DAF-7 could cooperate with signal transducer and activator of transcription (STAT) proteins, which can regulate nuclear gene expression with consequences for development and cell growth (Levy and Darnell, 2002; Wang and Levy, 2006). Consistent with a role of STAT in DAF-7 signaling, mutant *sta-1* resulted in alleviation of the germline development defect in the *xpc-1* background (Suppl. Figure 3F).

### DAF-7/TGF-beta signaling regulates the CEP-1/p53 induction in PGCs upon UV irradiation

We previously established that the *C. elegans* p53 orthologue, CEP-1, is induced in PGCs carrying persistent DNA damage and mediates their cell cycle arrest (Ou et al., 2019). The induction of CEP-1/p53 in PGCs is induced via FGF-like signaling emanating from IFE-4 induction in SGPs. Hence, we wondered whether ASI neuron signaling affects the CEP-1/p53 induction in PGCs. To this end, we used the established immunofluorescence (IF) staining method for PGCs in L1 larvae (Ou and Schumacher, 2021) and assessed the expression levels of CEP-1/p53 upon UV-induced DNA damage in the *xpc-1* mutant background. We used the PGL-3 antibody to label specifically PGCs (Figure 4A), and DAPI to normalize the CEP-1/p53 signal against DNA content, which we previously extensively validated as accurate IF signal normalization (Ou et al., 2019).

**Figure 4:**
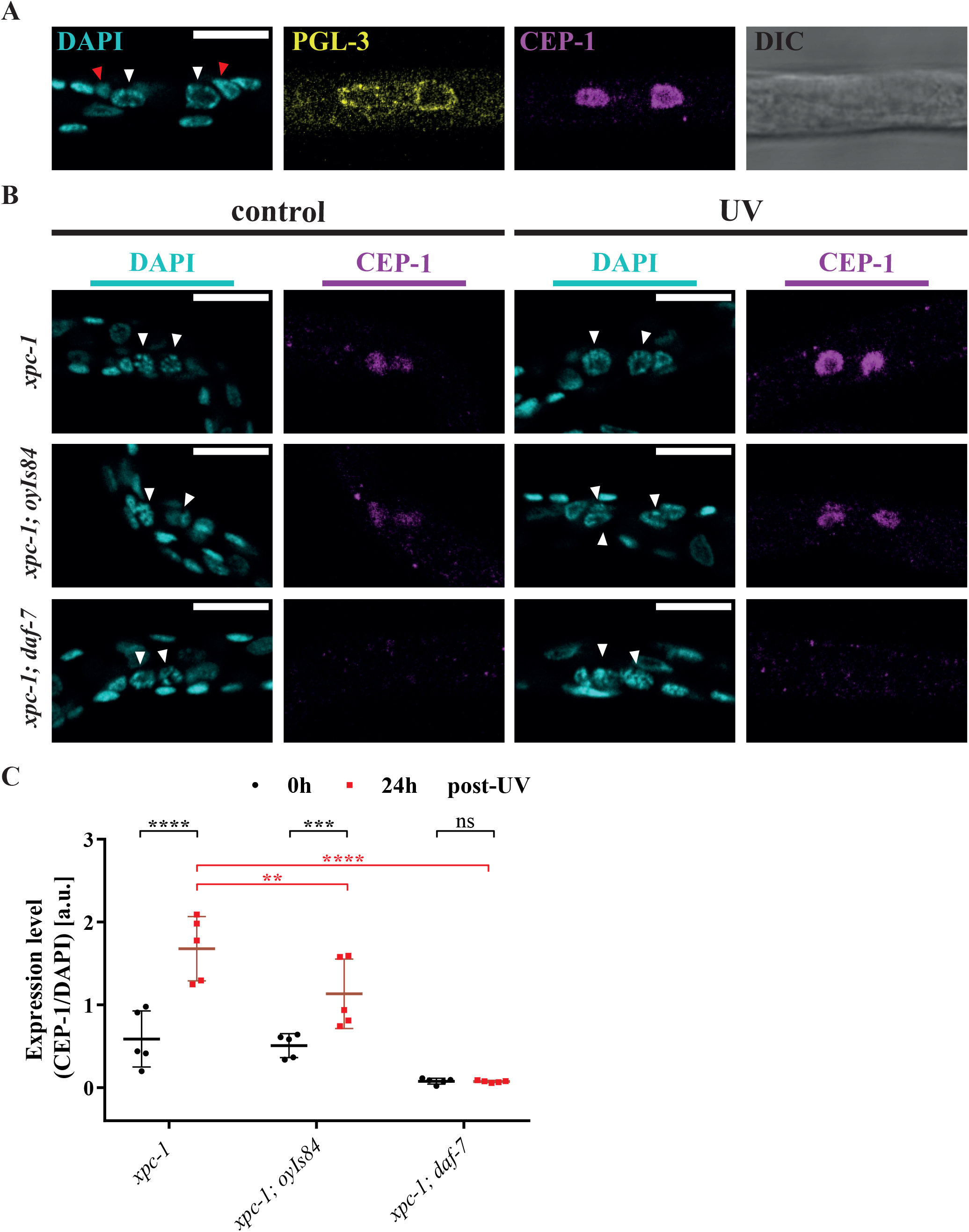
ASI neurons and DAF-7/TGF-β signaling regulate CEP-1/p53 induction upon DNA damage in PGCs. **(A)** Representative images of CEP-1/p53 expression levels in PGCs, which were identified by PGL-3 staining (24 h post-UV irradiation and incubation at 15 °C) in *xpc-1* mutant strain; white arrow heads indicate PGCs; red arrow heads indicate SGPs; scale bar represents 10 μm **(B)** Representative images of CEP-1 expression in *xpc-1, xpc-1; oyIs84* ASI ablated and *xpc-1; daf-7* TGF-β defective strains showing reduced CEP-1/p53 induction in ASI ablated animals, while, *daf-7* mutation results in total loss of CEP-1/p53 staining; scale bars represent 10 μm; white arrow heads indicate PGCs. **(C)** Quantification of CEP-1/p53 expression level as CEP-1/DAPI ratio in arbitrary units; 5 independent experiments were summarized and shown as mean ± SD; RM two-way ANOVA with matched values and Sidak’s multiple comparison test was performed; ns = not significant, **p<0.01, ***p<0.001, ****p<0.0001.

CEP-1/p53 protein levels in *xpc-1* mutants were induced in PGCs upon UV treatment, while *xpc-1; oyIs84* worms with ablated ASI neurons showed significantly reduced CEP-1/p53 levels. In *xpc-1; daf-7* mutants CEP-1/p53 protein was undetectable (Figure 4B and C), similarly to a *xpc-1; cep-1; oyIs84* control strain that carries a deletion mutation in *cep-1*. Control worms that were not UV-irradiated showed comparable basal levels of CEP-1 expression (Figure 4B and Suppl. Figure 5A). The complete absence of CEP-1/p53 protein in PGCs of *daf-7* mutants suggests that DAF-7 signaling is required even for baseline levels of CEP-1/p53 protein that might be established already prior to the ASI ablation in *oyIs84* transgenic animals.

Taken together, these results indicate that ASI neurons regulate, via secretion of DAF-7, the induction of CEP-1/p53 in PGCs upon persistent DNA damage.

### ASI neurons regulate the inheritance of DNA damage-induced mutations in PGCs

In humans, p53 suppresses tumorigenesis by preventing mutations resulting from DNA damage to expand through tumor cell proliferation. We next wondered whether failure to arrest genomically damaged PGCs through CEP-1/p53 could result in the expansion of mutations in germ cells and their subsequent inheritance. Given that not only CEP-1/p53 in PGCs but also the somatic niche cells (SGPs), and the ASI neurons regulate the arrest of PGCs, we could determine whether not only the germ cell DDR itself but also the regulatory input from the soma could impact the occurrence and inheritance of mutations. To assess the *de novo* mutational burden emanating from a dysregulated DDR in PGCs, we performed single worm whole genome sequencing (sw-WGS) of pairs of irradiated P0 and their direct F1 progeny. This comparison would allow us to selectively identify individual mutations that occurred in the germ cells originating from GG-NER deficient PGCs. Synchronized L1 worms were UV-irradiated and grown adults carrying developed germlines were singled onto plates with UVC-killed bacteria to perform egg laying before they were individually collected to provide P0 genome sequences. The respective eggs were incubated until the worms reached again young adult stage and picked as F1 generation. Both generations were subject to sw-WGS (Figure 5A). Genome sequences of *xpc-1* single mutant F1 worms were unattainable as they were not able to develop a germline amid UV treatment.

**Figure 5:**
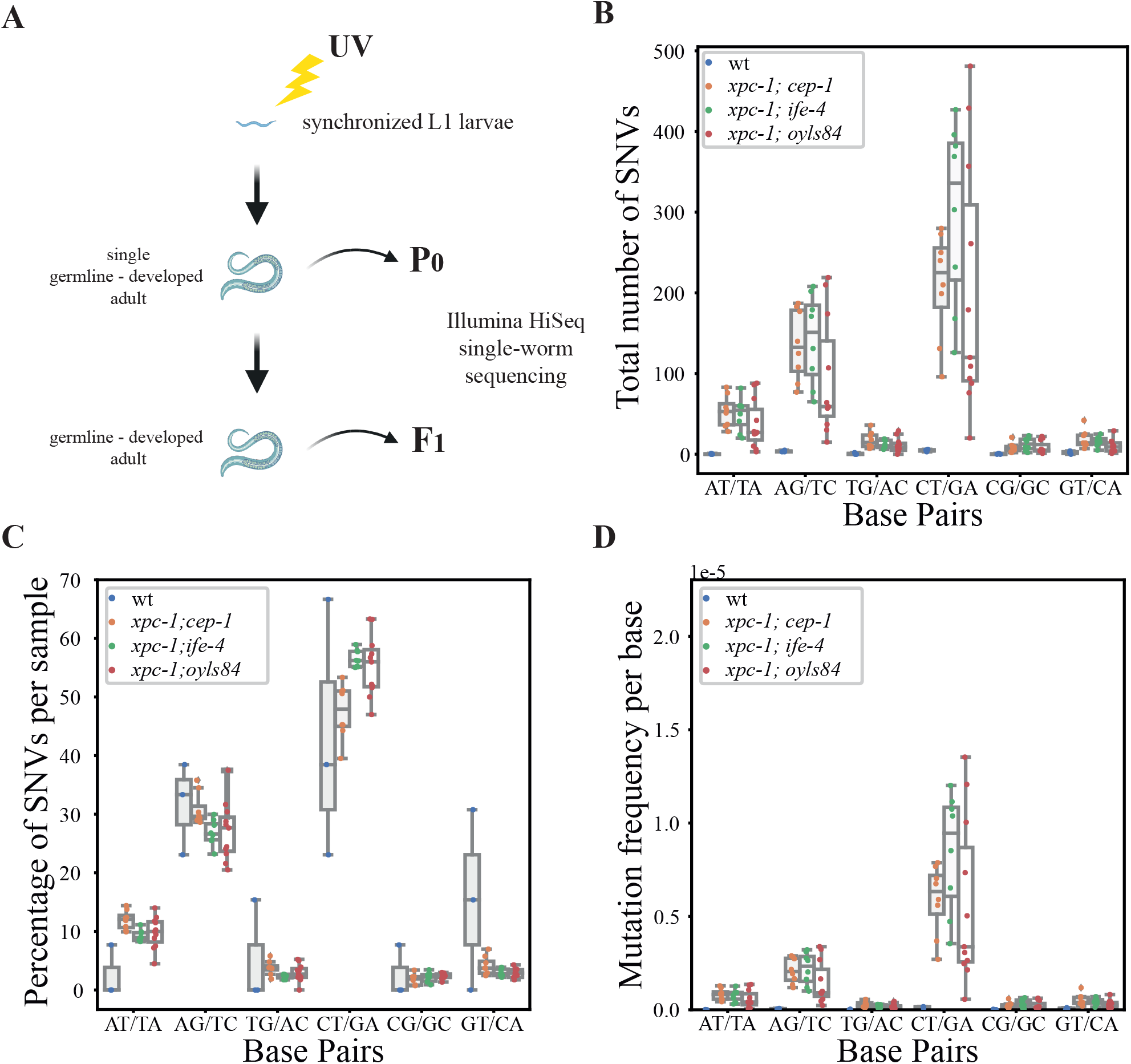
Loss of somatic niche or ASI neuron control of DDR in PGCs leads to inheritance of *de novo* mutations. **(A)** Experimental setup: UV-irradiated synchronized animals were scored for developed germlines and individually collected after successful egg-laying as P0 generation and direct offspring individuals as their F1 generation. The whole genomes of thus obtained pairs of single worms were sequenced. **(B)** Total numbers of identified SNVs divided into 6 different mutation classes, e.g., AT/TA depicts how many A>T, respective T>A mutations occurred. Color-coded are the 4 different strains. Each dot in each of the 6 groups represents the SNV calling results of one single worm WGS F1 generation sample minus SNVs that were found in the P0 generation of the corresponding strain. **(C)** Percentage of SNVs per sample. For each sample the sum of the percentages of the 6 groups add up to 100%. All strains show the highest increase in C>T/G>A mutations, a common pattern of UV-induced lesions converted into genomic substitutions. **(D)** Mutational frequency per base. The number of mutations that were reference A or T were divided by the total AT content of the ce11 genome (64745154 bp), the reference G or C mutations were divided by the total GC content of the genome (35541247 bp). The y-axis depicts the resulting frequency per base in the scale 1e-5. Noticeable all 3 double mutant strains show a drastic increase in mutational frequency, especially for C>T/G>A.

The sequencing analysis of *de novo* mutations arising in F1 animals of irradiated P0 revealed several orders of magnitude increased mutational burden in *xpc-1; cep-1*, *xpc-1; ife-4*, and *xpc-1; oyIs84* double mutants compared to F1 of UV treated WT animals. Among the single nucleotide variants (SNVs) particularly C>T/G>A transition mutations were strongly enriched (with median *de novo* SNV counts for C>T/G>A transitions of 225 in *xpc-1; cep-1, 336 in xpc-1; ife-4, and 119 in xpc-1; oyIs84*), resembling the common pattern of UV-induced lesions converted to genomic substitutions (Volkova et al., 2020). On the contrary, WT worms only showed a small number of *de novo* mutations consistent with the accurate repair of UV-induced lesions by GG-NER (with a maximum median *de novo* SNV count of 5 in C>T/G>A transitions) (Figure 5B). All 6 base pair SNV classes were significantly enriched in the 3 mutants compared to WT F1 genomes after multiple test correction (Supplement Table 1). Closer inspection of those SNVs revealed that the distribution and type of mutations were similar in all samples, indicating that even the few mutations detected in WT are similar to those in the double mutants (Figure 5C). Indeed, no comparison between WT and the 3 mutants showed significant differences in the type of *de novo* mutations (Supplement Table 1). Consistent with the increased number of total SNVs, also the mutation frequency per base was strongly increased when either CEP-1/p53 acting in the PGCs, IFE-4 acting in the somatic SGP niche cells, or the ASI neurons were defective (Figure 5D). Similar to the statistics in Figure 5B, all 6 base pair SNV classes were significantly elevated in the 3 mutants compared to WT F1 genomes and after multiple test correction (Supplement Table 1).

Taken together, our results show that the somatic niche and sensory ASI neurons safeguard the genomic integrity of PGCs and thus determine the inheritance of mutations occurring in germ cells.

## Discussion

To preserve the inheritance of stable genomes, organisms evolved specialized DNA repair and DNA damage response mechanisms, including the highly conserved GG-NER and p53 pathways, respectively. Genome integrity of PGCs, which are the origin of all gametes, is of utmost importance to maintain stable genomes through the generations. Here, we shed light on the non-cell-autonomous regulation through which the soma impacts the control of genome integrity in PGCs. We determined that the non-cell-autonomous DDR in the genomically compromised germline primordium is regulated by the signaling of environmental cue-sensing ASI neurons and the downstream TGF-beta signaling pathway. These data suggest that the nervous system transmits environmental information to regulate the DDR in PGCs and impacts the inheritance of mutations throughout generations (Figure 6).

**Figure 6:**
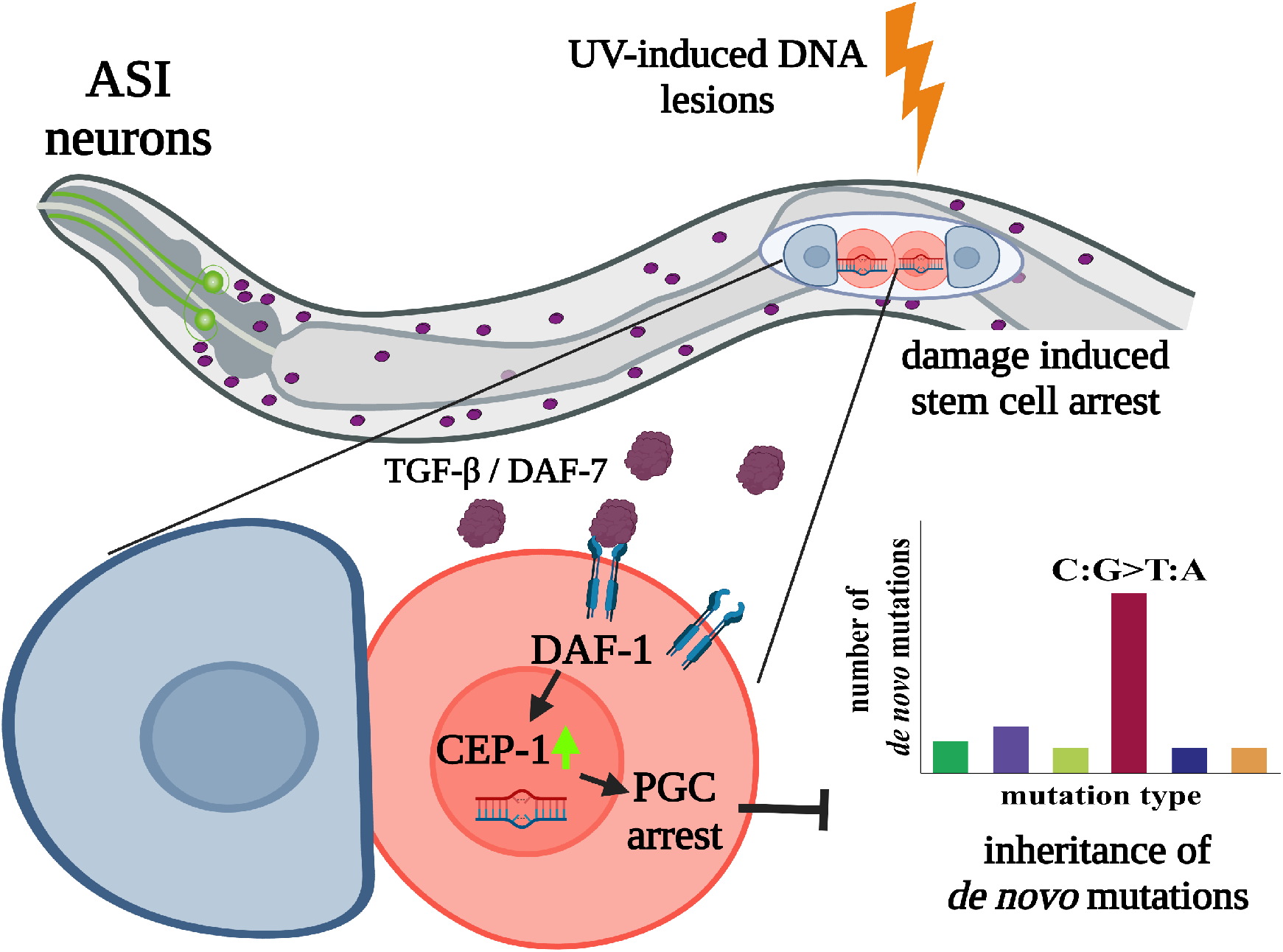
Neuronal signaling regulates the DNA damage response in primordial germlines and suppresses inheritance of *de novo* mutations. Release of TGF-beta ligand DAF-7 by amphid ASI neurons and reception via TGF-beta receptor 1 DAF-1 in PGCs is required for CEP-1-dependent DDR in the primordial germline upon persistent DNA damage and for thus suppressing the inheritance of *de novo* germline mutations.

We previously demonstrated a crosstalk between PGCs and its somatic niche that is comprised of the two SGPs (Ou et al., 2019). Likewise, the apoptotic response to DNA damage in meiotic pachytene cells during oogenesis was shown to be subject to regulation by the ASJ neurons (Sendoel et al., 2010) and by intestinal PMK-1/p38 signaling (Soltanmohammadi et al., 2022). Vice versa, the DDR in germ cells also influences the soma such as elevating somatic endurance amid unrepaired meiotic DNA damage (Ermolaeva et al., 2013; Soltanmohammadi et al., 2022). Furthermore, endocrine signals have been observed to non-cell-autonomously influence stem cell proliferation and the DDR in mammals and *C. elegans* (Cinat et al., 2021; Gronke et al., 2019; Robinson-Thiewes et al., 2021; Xuefeng Chen, et al, 2011; Zhang et al., 2019). Somatic signaling cues are important for germline development and determine epigenetic memories that can last through several generations. It was recently established that neuronal signals can have intergenerational influence on developmental timings (Perez et al., 2021). Here, we determined that a specific pair of neurons, the ASI neurons, regulate the DDR in PGCs. Mechanistically, we establish that TGF-beta, that is exclusively produced in ASI neurons, is perceived by the DAF-1 TGF-beta receptor in the PGCs, where it is required for the induction of the central DDR regulator CEP-1/p53.

Here, we investigated the role of AMP-activated kinases within non-cell-autonomous DDR in germline primordium upon persistent UV-induced DNA lesions. KIN-29, which was shown via RNAi knockdown to be involved in germline development arrest, is known to have similarity to AMPKs and the ability to act non-cell-autonomously in sensory signaling affecting body size and dauer decision (Lanjuin and Sengupta, 2002). Furthermore, KIN-29 was shown to be under translational control of IFE-4 (Dinkova et al., 2005). These observations prompted us to assay the ability of KIN-29 and an additional independent AMPK subunit, AAK-2, to suppress the germline development defect in the *xpc-1* mutant background upon UV irradiation. Both mutants showed suppression as expected and guided us towards nutritional signals. AMPKs function in interpreting nutritional and environmental states (Hardie et al., 1998), which are initially sensed via chemosensory neurons. ASI neurons are important sensory neurons for the interpretation of nutritional, environmental and population-related cues (Bargmann and Horvitz, 1991; Bishop and Guarente, 2007; Golden and Riddle, 1984; Schackwitz et al., 1996). We used multiple approaches to inactivate ASI signaling and all resulted in an alleviated germline development defect amid persistent DNA damage in PGCs. ASI neurons are exclusively producing and releasing the TGF-beta ligand DAF-7 via dense-core vesicles. Indeed, the PGC arrest and subsequent germline development defect upon DNA damage critically depended on DAF-7 signaling. The TGF-beta signaling is moreover directly received in the PGC through the DAF-1 TGF-beta receptor as evidenced by the similar suppression of the germline development defect in *xpc-1* mutants by *daf-7* and a null mutation in *daf-1*. In contrast, the TGF-beta receptor 2/ *daf-4* mutant did not phenocopy its heteromeric-counterpart (Wrighton et al., 2008). Why DAF-4 did not show the same drastic effect as its potential phosphorylation target and binding-partner DAF-1 is unclear, since the allele *daf-4(e1364)* is described to have lost kinase activity (Gunther et al., 2000). DAF-4 was originally reported to bind bone morphology proteins (BMPs) as TGF-beta ligand and, therefore, was proposed as orthologue of the human TGF-beta receptor 2, the first binding receptor for TGF-beta ligands in mammalian systems (Estevez et al., 1993; Georgi et al., 1990). Whether DAF-7 as alternative TGF-beta ligand could bind preferably DAF-1 independently from DAF-4 was indeed suggested before (Gunther et al., 2000), and is consistent with the different binding affinities of distinct ligands to specific receptors (Derynck and Budi, 2019).

TGF-beta signaling was shown to regulate numerous (i) developmental programs, such as body size and reproductive aging, (ii) stress responses, like pathogen avoidance and autophagy, and (iii) the starvation response and dauer formation (Dineen and Gaudet, 2014; Luo et al., 2010; Moore et al., 2019; Ren et al., 1996; Xuefeng Chen, et al, 2011; Zhang et al., 2019).

In mammals, TGF-beta is a pleiotropic cytokine that interacts with several stress response factors either via its downstream Smad-signaling in epithelial cells, in mutual combination with ROS or even in crosstalk with p53 (Dupont et al., 2004; Kang et al., 2003; Krstić et al., 2015). TGF-beta was described to co-regulate gene transcription by p53 via downstream Smad-p53 complex formation and, similar to p53, TGF-beta is further known to regulate microRNA biogenesis in mammals promoting cell proliferation, which stands in direct contrast to commonly known tumor suppressive roles (Braun et al., 2008; Davis et al., 2010; Suzuki, 2018; Suzuki and Miyazono, 2010). The involvement of TGF-beta in the p53 regulation suggests that the influence of DAF-7 signaling in the non-cell-autonomous DDR is thus highly conserved between nematodes and mammals. Indeed, CEP-1/p53 protein in PGCs carrying persistent DNA damage was virtually undetectable in *xpc-1; daf-7* double mutants indicating that neuronally secreted TGF-beta is critical for CEP-1/p53 induction in PGCs.

Importantly, we determined that the regulation of the CEP-1/p53-mediated DDR in PGCs by either IFE-4 acting in the somatic niche cells or by ASI neurons impacts the inheritance of mutations. Thus far somatic influences on subsequent generations have been confined to epigenetic effects (notably small RNA-mediated effects) that would last through several generations but are not manifested in a species’ gene pool. In contrast, we determined that defects in neuronal control of the PGCs’ DDR could result in inheritance of *de novo* mutations through a defective CEP-1/p53 controlled PGC arrest (Figure 6). Our results suggest that a single pair of sensory neurons that transmit environmental signals to the physiological states of the animal, determine whether mutations are transmitted and could become part of the gene pool. This finding further challenges the Weismann barrier concept, which pertains that somatic influences are blocked from impacting the heritable information and instead suggests that the state of sensory neuron signaling determines genetic inheritance.

## Acknowledgements

We thank S. Strome for sharing the anti-PGL-3 antibody. Worm strains were provided by the National Bioresource Project (supported by The Ministry of Education, Culture, Sports, Science and Technology, Japan), the Caenorhabditis Genetics Center (funded by the NIH National Center for Research Resources, US), and the *C. elegans* Gene Knockout Project at the Oklahoma Medical Research Foundation (part of the International *C. elegans* Gene Knockout Consortium). Schemes in Figure 3A and 5A were created with BioRender.com. B.S. acknowledges funding from the Deutsche Forschungsgemeinschaft (SCHU 2494/3-1, SCHU 2494/7-1, SCHU 2494/10-1, SCHU 2494/11-1, CECAD EXC 2030 – 390661388, SFB 829, KFO 286, KFO 329, GRK 2407, and SCHU 2494/15-1 to B.S. and O.R.), the Deutsche Krebshilfe (70112899), the H2020-MSCA-ITN-2018 (Healthage and ADDRESS ITNs) and the John Templeton Foundation Grant (61734) to B.S. and O.R.

B.S is co-founder of Agevio Therapeutics, Inc. The authors declare no other competing interests.

## Author contributions

S.U. performed all *C. elegans* experiments and analyzed the data, D.H.M analyzed single worm sequencing data, B.S. coordinated the project and together with S.U. and O.R. designed the study. All authors wrote the paper.

## Material and Methods

### *C. elegans* strains

Strains used in this study were derived from N2 Bristol wild type. Unless otherwise indicated. Worms were grown under standard laboratory conditions and fed with OP50 *Escherichia coli* on nematode growth medium (NGM) agar. Strains used for this study are as following: N2 (Bristol) as wild type, BJS355 *xpc-1(tm3886)*, FX00684 *ife-4(tm684)*, XY1054 *cep-1(lg12501)*, BJS251 *xpc-1(tm3886); ife-4(tm684)*, BJS68 *xpc-1(tm3886); cep-1(lg12501)*, VC539 *kin-29(gk270)*, BJS761 *xpc-1(tm3886); kin-29(gk270)*, RB754 *aak-2(ok524)*, BJS973 *xpc-1(tm3886); aak-2(ok524)*, PY7505 *oyIs84[gpa-4p::TU#813+gcy-27p::TU#814* + *gcy-27p::GFP* + *unc-122p::dsRed]*, BJS896 *xpc-1(tm3886); oyIs84[gpa-4p::TU#813* + *gcy-27p::TU#814* + *gcy-27p::GFP* + *unc-122p::dsRed]*, BJS261 *xpc-1(tm3886); bnIs1[pie-1p::GFP::pgl-1* + *unc-119*(+)], BJS993 *xpc-1(tm3886); oyIs84[gpa-4p::TU#813* + *gcy-27p::TU#814* + *gcy-27p::GFP* + *unc-122p::dsRed]; bnIs1[pie-1::GFP::pgl-1* + *unc-119*(+)], CF4126 *muEx641[pPC30(pgpa-4::GFP::Tetx)* + *punc-122::RFP]*, BJS990 *xpc-1(tm3886); muEx641 [pPC30(pgpa-4::GFP::Tetx)* + *punc-122::RFP]*, SP1735 *dyf-7(m537)*, BJS983 *xpc-1(tm3886); dyf-7(m537)*, RB712 *daf-18(ok480)*, BJS907 *xpc-1(tm3886); daf-18(ok480)*, PHX4372 *daf-7::V5(syb4362)*, BJS989 *xpc-1(tm3886); ksIs2 [daf-7p::GFP* + *rol-6(su1006)]*, CB1372 *daf-7(e1372)*, MT1438 *daf-1(n690)*, DR40 *daf-1(m40)*, CB1364 *daf-4(e1364)*, BJS969 *xpc-1(tm3886); daf-1(n690)*, BJS970 *xpc-1(tm3886); daf-1(m40)*, BJS968 *xpc-1(tm3886); daf-4(e1364)*, BJS974 *xpc-1(tm3886); daf-7(e1372)*, BJS359 *xpc-1(tm3886); rrf-1(pk1417)*, BJS583 *xpc-1(tm3886); rde-1(ne219); qyIs102[fos-1ap::rde-1* + *myo-2p::YFP* + *unc-119*(+)], BJS994 *xpc-1(tm3886); cep-1(lg12501); oyIs84[gpa-4p::TU#813* + *gcy-27p::TU#814* + *gcy-27p::GFP* + *unc-122p::dsRed]*, BJS995 *xpc-1 (tm3886); sta-1(ok587)*, BJS1027 *xpc-1(tm3886); daf-7::V5(syb4362)*.

All experiments were conducted at 20 °C except experiments involving mutants with temperature sensitive abnormal dauer formation (i.e. *daf* mutants) or ASI defective strains that were conducted at 15 °C to avoid dauer formation.

### PHX4372 *daf-7::V5(syb4362)* generation

The PHX4372 *daf-7::V5(syb4362)* was purchased from SunyBiotech. To attain a DAF-7::V5 fusion protein, a CRISPR-Cas mediated precise sequence knock-in of the V5 tag into the endogenous locus of *daf-7* (B0412.2.1) before the termination codon of wild type N2 strain was generated. The PHX4372_strain was genotyped and backcrossed in our lab before crossed into the GG-NER deficient *xpc-1* background.

### Germline development assay

Worms were synchronized via 3 hours of time-restricted egg-laying on plates seeded with *E. coli* as food source. Respectively, eggs were incubated for 12 – 14 hours at 20 °C or 14 - 16 hours at 15 °C. During this time, hatching progress was monitored to mock- or UVB-irradiate when all L1 larvae hatched. When somatic development was completed three or four days later, respectively, worms were examined utilizing the stereomicroscope (Leica M80) and the percentage of worms with developed germlines was scored.

### Primordial germ cell (PGC) proliferation assay

Strains carrying the transgene *bnIs1*[P_*pie-1*_*::GFP::pgl-1* + *unc-119*(+)] were bleach-synchronized and L1 larvae were subjected to UVB irradiation or mock-treatment before feeding and incubated at 15 °C. For the kinetic assays, the number of germ cells was scored every 2-4 hours after expected proliferation onset post UVB irradiation with fluorescence microscope. For PGC proliferation assays the initial number of germ cells was scored at 0 h.

### RNAi knockdown

For the RNAi experiments, strains of *E. coli* HT115 (DE3) transformed with isopropyl-β-D-thiogalactopyranosid (IPTG)-inducible vectors that encode double-stranded RNA against target genes were thawed from the *C. elegans* RNAi feeding library constructed by Julie Ahringer’s group at The Wellcome CRC Institute (University of Cambridge, Cambridge, UK) (obtained via Source BioScience with the product code 3318_Cel_RNAi_complete) or the *C. elegans* ORFeome-RNAi v1.1 library generated by Prof. Marc Vidal’s group at Center for Cancer Systems Biology and Department of Cancer Biology, Dana-Farber Cancer Institute and Department of Genetics, (Harvard Medical School, Boston, USA) (obtained via Geneservice (now as Source BioScience) with the product code 3320_Cel_ORF_RNAi). *E. coli* feeding strain HT115 (DE3) with corresponding RNAi was cultured using liquid LB medium containing ampicillin (100 mg/mL) over-night at 37 °C. Over-night cultures were inoculated at OD_600_ 0.25 in fresh LB medium containing ampicillin (100 mg/mL) at 37 °C and after 90 min RNAi production was induced by addition of 1 mM IPTG for 4 hours before seeded on RNAi agar plates. Subsequently, seeded plates were incubated over-night at 37°C. For RNAi feeding, worms of each strain were synchronized at L4 larval stage as P0 generation when transferred to RNAi agar plates that contain standard NGM agar, Ampicillin (100 mg/mL), IPTG (1mM), and *E. coli* feeding strain HT115 (DE3) with the corresponding RNAi. Worms were incubated at 15 °C. Every four days the L4 larvae of the next generation were transferred to fresh RNAi plates to maintain the RNAi knockdown efficiency until performing further experiments.

### UV irradiation

Worms were irradiated with 310 nm UVB light using Philips UV6 bulbs in a Waldmann UV236B irradiation device. UVX digital radiometer and UVX-31 probe from UVP was used prior to every irradiation to calculate irradiance. Dosages of 25 mJ/cm^2^ were used in experiments, except additionally specified.

### Microscopy

For microscopy of worms, differential interference contrast (DIC) and fluorescence images for quantification were taken using Zeiss Axio Imager. M1 microscope while the evaluation of germline development phenotype was carried out using Leica M80 stereomicroscope. For the representative images with high resolution and IF staining, the Zeiss Meta 710 confocal laser scanning microscope was used.

### Immunofluorescence staining

Worms of indicated strains were synchronized by bleaching at L1 larval stage and subject to UVB irradiation or mock-treated on NGM plates without OP50 *E. coli*. Either worms were directly harvested for the staining procedure as 0 h control group or after 24 hours incubation time at 15 °C to measure DNA damage induced protein expression. Worms were washed 3 times with M9 buffer and centrifuged at 800 x g for 2 min to remove bacteria. Prior to the staining procedure, per strain and condition 5 μl fixing buffer (consisting of 27.5 mM HEPES pH7.4, 130 mM NaCl, 52.8 mM KCl, 2.2 mM EDTA, 550 mM EDGA, 0.1 % Tween 20, and 3 % Paraformaldehyde) was prepared and aliquoted to 0.5-ml tubes and a box was filled with dry ice. 5 μl of concentrated worms was transferred to the fixing buffer and mixed by pipetting up and down 10 times before further transferred as 2 μl droplet to self-made Poly L-Lysine coated Diagnostic Microscope Slide (ThermoScientific, ER-303B-CE24). A 24 x 24 mm coverslip was then attached to the transferred drop and the slide was left at room temperature for 2 minutes of incubation with slight tapping on top of the coverslip. The slide was put into the box with dry ice and kept for at least 30 minutes until performing freeze-cracking procedure. Once the freeze-cracking was done, slides were immediately transferred to prechilled ice-cold methanol-filled chamber. After 10 minutes of incubation, slides were washed 3 times with PBST (dissolving 240 mg KH_2_PO_4_, 1.44 g NaHPO_4_, 8 g NaCl, 200 mg KCl, and 1 ml of Tween 20 in 1 L of distilled H_2_O) for 10 minutes each time. For blocking, slides were incubated with 10 % donkey serum diluted in PBST for 30 min at room temperature in humid chambers. The primary antibodies were diluted in 10% donkey serum with PBST in various dilution as described in (H. L. Ou & Schumacher, 2021) (rabbit anti-V5 1:50 [Abcam, ab206566], goat anti-CEP-1 antibody, 1:200 dilution; rat anti-PGL-3, 1:10000 dilution (PGL-3 antibody was generously provided by Prof. Susan Strome, University of California Santa Cruz, USA) and the slides were incubated in the humid chamber at 4°C overnight. The slides were washed 3 times with PBST for 10 minutes each time, before incubated with secondary antibodies: donkey anti-goat IgG AlexaFluor 488 or AlexaFluor 647 (Jackson Immuno Research #705-545-003 and #705-605-147); donkey anti-rat IgG Alexa 488 or Alexa 594 (Invitrogen #A21208 and #A21209); donkey anti-rabbit IgG Alexa 594 (Invitrogen #A21207); all in 1:350 dilution with incubation at room temperature for 2 hours. After 3 times washing with PBST, slides were mounted with DAPI Fluoromount-G mounting medium (Southern Biotech) and sealed with nail polish when the mounting medium was dry. The slides were then ready for microscopy imaging.

### Quantification of immunofluorescence images

The worms with immunofluorescence staining were imaged with fixed exposure time in order to compare between different strains and treatments. In CEP-1 immunofluorescence staining assay, PGCs were manually selected via PGL-3 staining and the fluorescence intensity of both CEP-1 and DAPI were measured with ImageJ software (Rasband, W.S., ImageJ, U. S. National Institutes of Health, Bethesda, Maryland, USA, http://imagej.nih.gov/ij/, 1997-2015). For each PGC set the respective unspecific background was subtracted and ratios were calculated using CEP-1 fluorescence intensity normalized with corresponding DAPI intensity before plotted.

### DiI staining

Bleach synchronized L1 worms were transferred to OP50 E. coli seeded NGM plates. After 3 hours incubation at 20 °C, animals were washed off with M9 buffer into 15 ml Falcon tubes and washed 3 times to remove bacteria. Lipophilic DiI dye [2 mg/ml] stock was diluted 1:200 in M9 buffer and added to washed worms. After 1 hour of incubation at RT, light protected and rolling at 200 rpm, worms were spun down at 1200 rpm and washed 3 times. Worms were subsequently plated on seeded NGM plates for 1 hour of crawling to remove intestinal dye. Animals were picked onto 2 % agarose pads and mounted in levamisole for imaging.

### Preparation of single worm sequencings for mutational burden post UVB exposure

Bleach synchronized L1 worms were transferred to empty NGM plates, UVB irradiated and subsequently fed with OP50 *E. coli*. Animals were incubated at normal laboratory conditions until reaching YA stage. Double mutants *xpc-1; ife-4* and *xpc-1; cep-1* were incubated at 20 °C. Double mutant *xpc-1; oyIs84* was incubated at 15 °C. Small NGM-plates were freshly seeded with OP50 over-night culture, dried and subsequently bacteria were killed by 20 min UVC exposure. 10 individual worms scored for a partially/fully developed germline were singled onto UVC-OP50 plates, incubated until first eggs were laid and immediately picked into 10μl M9 buffer as P0 generation and flash frozen on dry ice. The laid eggs were incubated until next generation reached YA stage and were picked as paired F1 generation.

### Single worm DNA library preparation

Illumina DNA preparation and sequencing were performed by Cologne Center for Genomics (GGC, Cologne). Single worm lysis and Genomic DNA Tagmentation were performed according to manufacturer’s manual (Illumina DNA Prep Reference Guide). Amplification of Tagmented DNA was done with 12 PCR Cycles. Sequencing was performed via HiSeq4000 2×76 bps for wt, *xpc-1; ife-4* and *xpc-1; cep-1* samples and via NovaSeq 2×151bp for *xpc-1; oyIs84* samples.

### Single worm *de novo* mutation analysis

The fastq files were trimmed and preprocessed using Fastp v0.2 (Chen et al., 2018). The remaining reads were then aligned to the *C. elegans* reference genome (ce11) with BWA v0.7.17 (parameter setting: bwa mem -M -t 8 -K 100000000 ce11.fasta) (Li, 2013). The mapped files were converted to BAM and sorted with samtools v1.6 (Li et al., 2009). After sorting, duplicated reads were marked with GATK-4.1.0.0’s MarkDuplicates (Van der Auwera and O’Connor, 2020). All samples had comparable numbers of mapped reads (~68.5M +-13.5M).

Mutations were called with Strelka-2.9.10 (Kim et al., 2018) on single worms in the F1 generation and on merged cohorts for each strain in the P0 generation. The resulting VCF-files were filtered for SNVs that passed the quality filter, i.e., SNVs that contained “PASS” in the 7th column of the VCF-file.

To further reduce the number of false positives we filtered the VCF-files of the F1 generation with a minimum GQX value of 30 and a maximum homopolymer stretch of 6. SNVs of the cohort samples were kept if at least half of the samples were not homozygous reference.

To obtain the SNVs that were newly introduced by the UV irradiation in the F1 generation we subtracted SNVs found in P0 from the SNV list in the F1 generation for each respective strain. The total number of F1 SNVs per sample was divided into the 6 different potential mutation pairs:

A->T mutation, respective T->A on the complementary strand
A->G, respective T->C
T->G, respective A->C
C->T, respective G->A
C->G, respective G->C
G->T, respective C->A

The mutation frequency per sample was calculated by dividing the number of mutations of each of the 6 groups by the total of all mutations in the respective sample to obtain values between 0 and 1, that add up to 1 for each sample.

The mutation frequency per base was calculated by dividing the number of mutations of each group by the GC, respective AT, content of the whole genome. The number of AT/TA, AG/TC, and TG/AC mutations was divided by the number of ATs in the ce11 genome (number of GCs=35541247, number of ATs=64745154, total genome size=100286401), and the number of CT/GA, CG/GC, and GT/CA mutations by the number of GCs.

### Statistical analysis

Statistical methods that were used and error bar descriptions are specified in the figure legends. Randomization was not applied because the group allocation was guided by the genotype of the respective mutant worms. Worms of a given genotype were randomly selected from large strain populations for each experiment without any preconditioning. Blinding was not applied as the experiments were carried out under highly standardized and predefined conditions such that an investigator-induced bias can be excluded. Statistical analyses were performed using GraphPad Prism 7 software package (GraphPad Software, La Jolla California USA, www.graphpad.com).

---

**Suppl. Figure 1: Mock-treatments and *xpc-1* germline development phenotype. (A, C)** Germline development of indicated *C. elegans* double mutants are not affected upon mock-treatment (0 mJ/cm^2^ UV); worms were scored side-by-side with tested double mutants from Figure 1. **(B, D)** Germline development of single mutants, used to generate double mutants in the *xpc-1* background for studies of AMPK KIN-29 **(B)** and AMPK AAK-2 **(D)**, are not affected upon mock- or UV-treatment. Ratio of worms with developed germlines scored 3 days post mock- or UV-irradiation. 3 independent experiments with each 3 technical replicates were summarized and are shown as mean ± SD. **(E)** Representative images of *xpc-1* mutants 3 days post mock- or UV-treatment. Majority of animals are lacking developed germlines. Few *xpc-1* animals, having either egg-less germlines or tumorous growth in germline area (highlighted with white line), were nevertheless scored as worms with developed germline (~ 5%); percentages vary among experiments and represent a rough estimation; asterisks indicate vulva position; scale bars represent 50 μm.

**Suppl. Figure 2 Lipophilic DiI-staining confirms functional impairment of ASI neurons in *xpc-1; oyIs84* animals. (A)** In wild type (N2) animals functional ASI neurons can take up the lipophilic dye DiI. **(B)** Although ASI-specific GFP signal remains, no DiI-stain was taken up suggesting ASI inactivity in *xpc-1; oyIs84[gpa-4p::TU#813* + *gcy-27p::TU#814* + *gcy-27p::GFP* + *unc-122p::dsRed]* mutants; white arrow heads indicate ASI position; scale bars represent 20 μm. Animals were bleach-synchronized and DiI staining was performed within 4 hours post initial feeding.

**Suppl. Figure 3: DAF-18 is dispensable for germline development defect and control mutants have no effects on physiological germline development. (A)** ASI ablation had no influence on starvation induced PGC cell cycle arrest; PGC proliferation kinetics scored in animals carrying transgene *bnIs1*[P_*pie-1*_::GFP::*pgl-1* + *unc-119(+)]*. Animals were bleach synchronized, hatched in starved conditions and transferred to empty plates until PGCs were counted. 3 independent experiments are combined and shown as dot blot with mean ± SD. **(B)** Single mutants of *oyIs84[gpa-4p::TU#813* + *gcy-27p::TU#814* + *gcy-27p::GFP* + *unc-122p::dsRed]*, *dyf-7* and *daf-18* used to generate respective double mutants in *xpc-1* background, do not show any germline development defect phenotype upon UV. **(C)** Tumor suppressor PTEN mutant, *xpc-1; daf-18*, did not show any restoration of the germline development post treatment. **(D)** DAF-7 deficiency did not lead to spontaneous escape from PGC cell cycle arrest in L1 larvae, which hatched in nutrient deprived conditions (M9 buffer) prior to feeding. PGC number was counted from IF staining experiments at 0h control conditions. Summarized PGC numbers from 5 independent experiments are shown as dot blot with mean ± SD. **(E)** Single mutants of TGF-beta ligand *daf-7* and receptors *daf-1* and *daf-4* used to generate respective double mutants in *xpc-1* background, do not show any germline development defect phenotype upon UV. **(F)** STAT-kinase mutant strain *xpc-1; sta-1* showed significant alleviation of germline development defect post-UV irradiation. (B-C and E-F) Ratio of worms with developed germlines scored 4 days [incubation at 15 °C] (B,C,E) or 3 days [incubation at 20 °C] (F) post-UV irradiation. 3 independent experiments with each 3 technical replicates were summarized and shown as mean ± SD. One-way ANOVA with Dunnett’s multiple comparison test was performed; *p<0.05, **p<0.01, ****p<0.0001, ns=not significant.

**Suppl. Figure 4: ASI exclusive DAF-7 expression. (A)** *daf-7* promoter-driven GFP expression is exclusively located in ASI neurons. **(B)** DAF-7::V5 fusion protein is also exclusively located in ASI neurons while only dim background staining was observed in *xpc-1* control animals. Scale bars represent 20 μm.

**Suppl. Figure 5: CEP-1/p53 expression levels 0 h and 24 h post-UV of internal control worms. (A)** Representative images of CEP-1/p53 expression levels and quantification as CEP-1/DAPI ratio in arbitrary units in PGCs of wild type (wt) and *xpc-1; cep-1; oyIs84[gpa-4p::TU#813* + *gcy-27p::TU#814* + *gcy-27p::GFP* + *unc-122p::dsRed]* animals at 0 h control timepoint or 24 h post UV-irradiation from same analysis shown in Figure 4; wt shows moderate induction of CEP-1 expression 24 h post UV-irradiation; CEP-1 null strain *xpc-1; cep-1; oyIs84* was included to confirm staining specificity of the anti-CEP-1 antibody; scale bars represent 10 μm; white arrow heads indicate PGCs; each dot represents one of 5 independent experiments, which are shown as mean ± SD; RM two-way ANOVA with matched values and Sidak’s multiple comparison test was performed; ns = not significant, **p<0.01, ***p<0.001, ****p<0.0001.

**Supplement Table 1**. Summarized statistics of shown experiments sorted by appearance with detailed information about respectively used statistical methods.

**Supplement Table 2**. Overview of all identified single nucleotide polymorphisms (SNVs) in mutational cohorts and filtered *de novo* SNVs in respective F1 animals.

