## Supplementary figures and images for "Sensory neurons safeguard from mutational inheritance by controlling the CEP-1/p53-mediated DNA damage response in primordial germ cells"

### Supplementary Figures 1-5

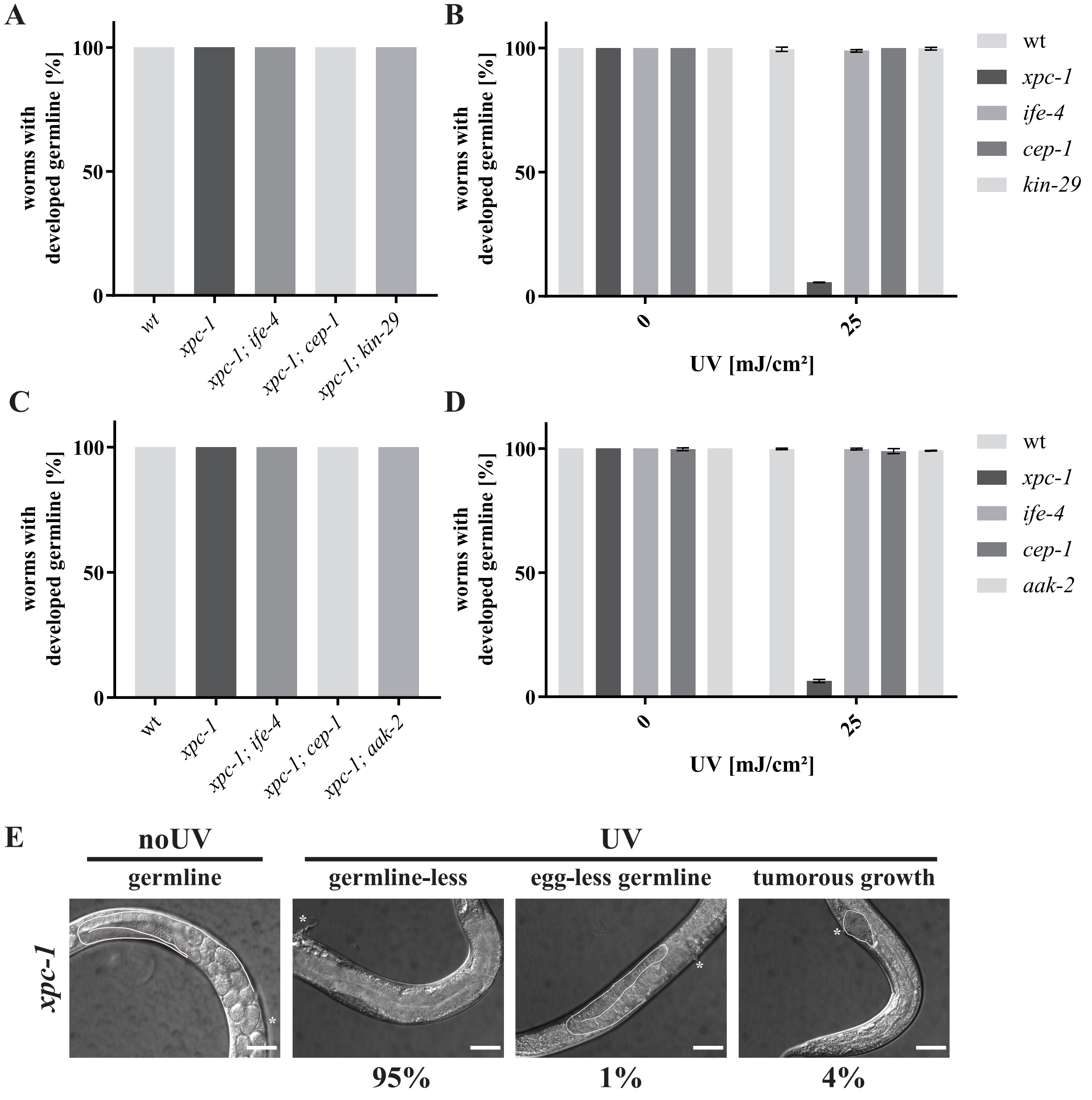

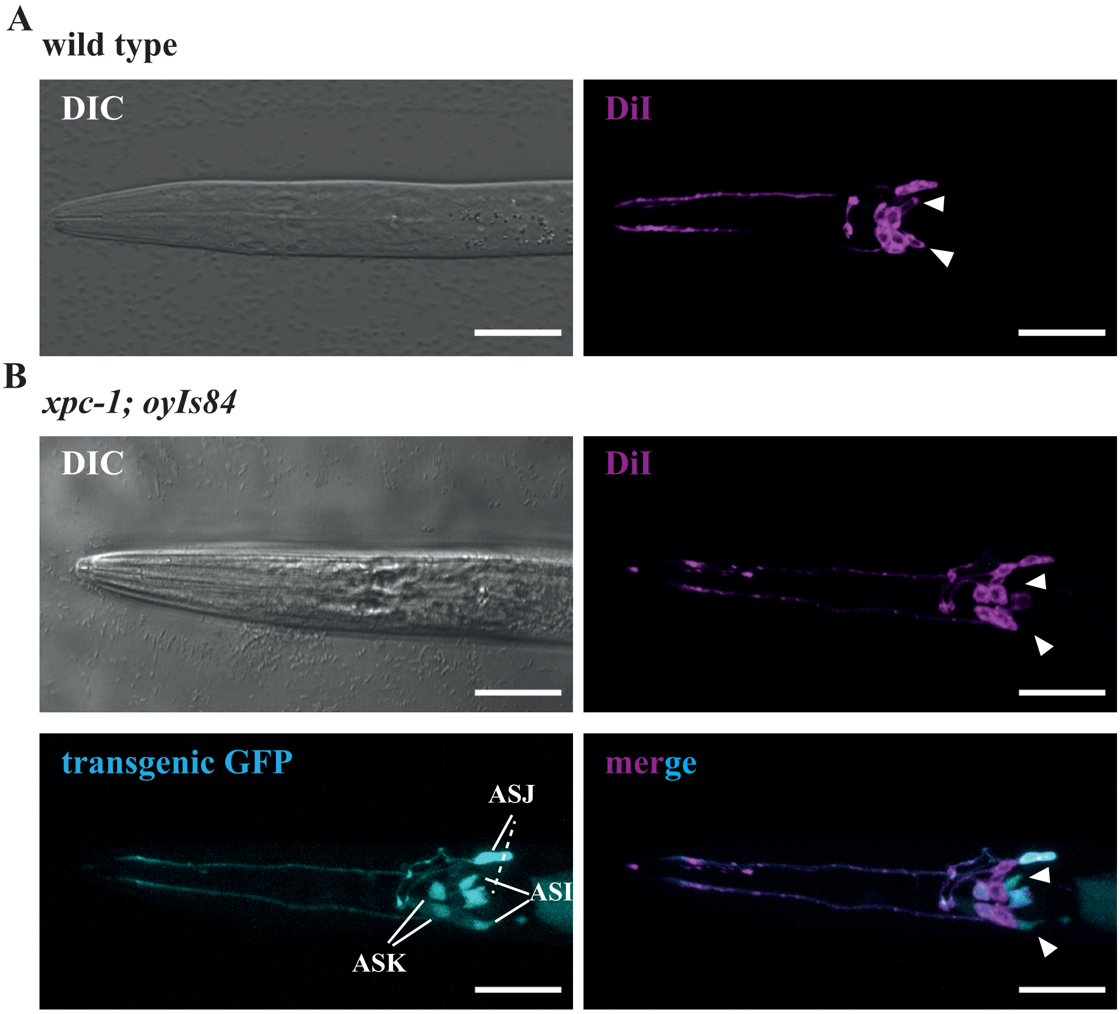

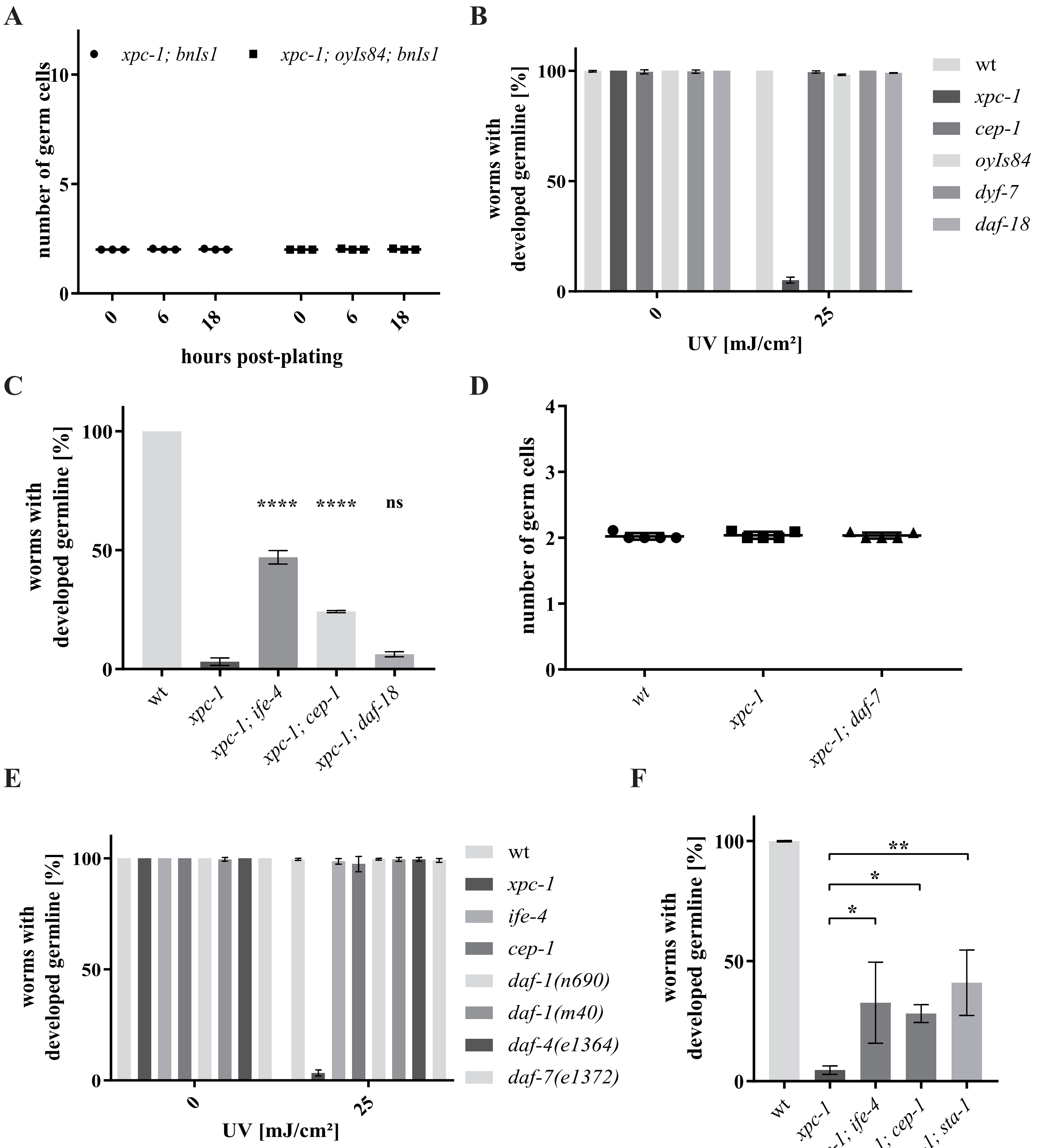

**Suppl.Fig.4**

**A** *xpc-1; ksIs2 [daf-7p::GFP + rol-6(su1006)]*

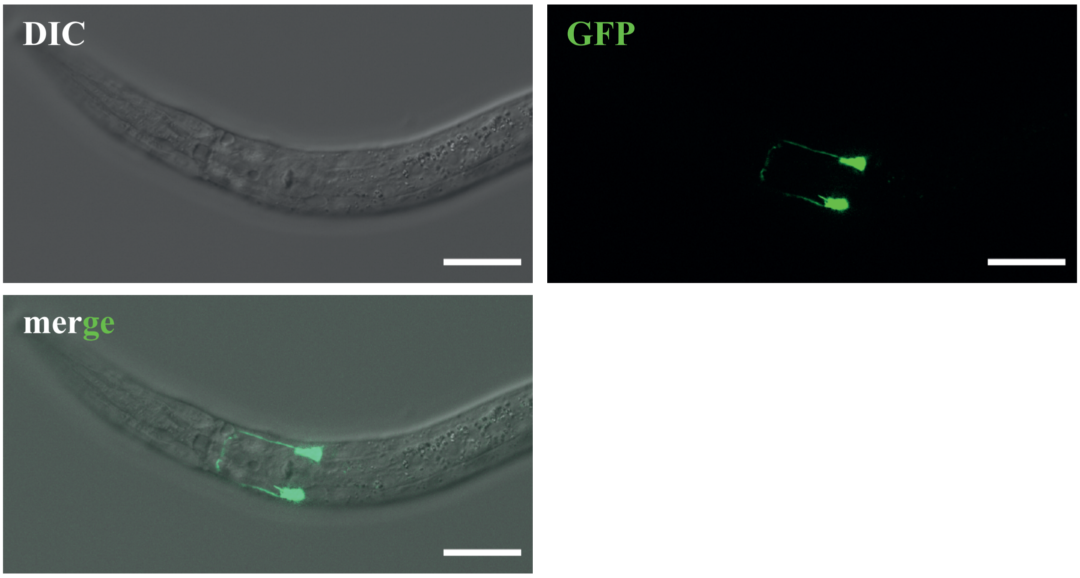


**B** *xpc-1; daf-7::V5*

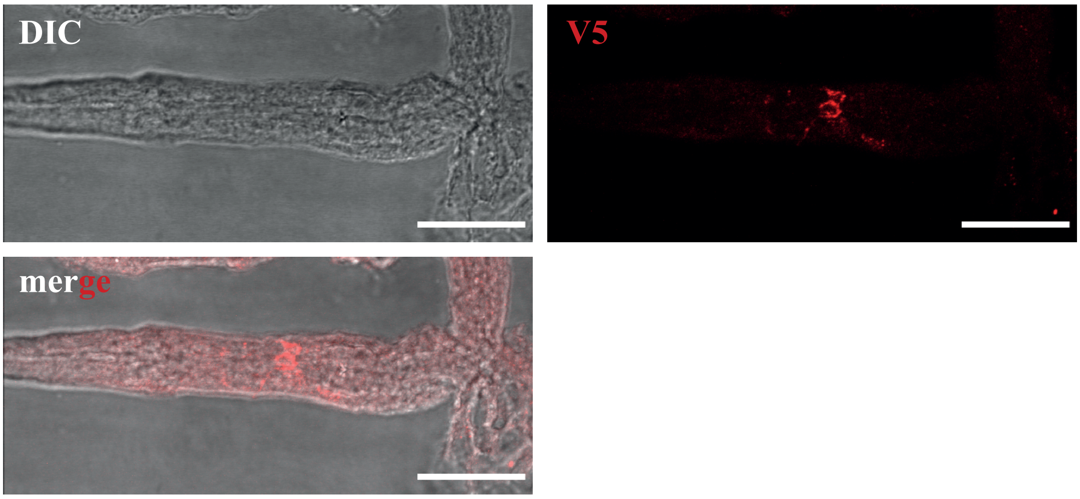


*xpc-1* (background control)

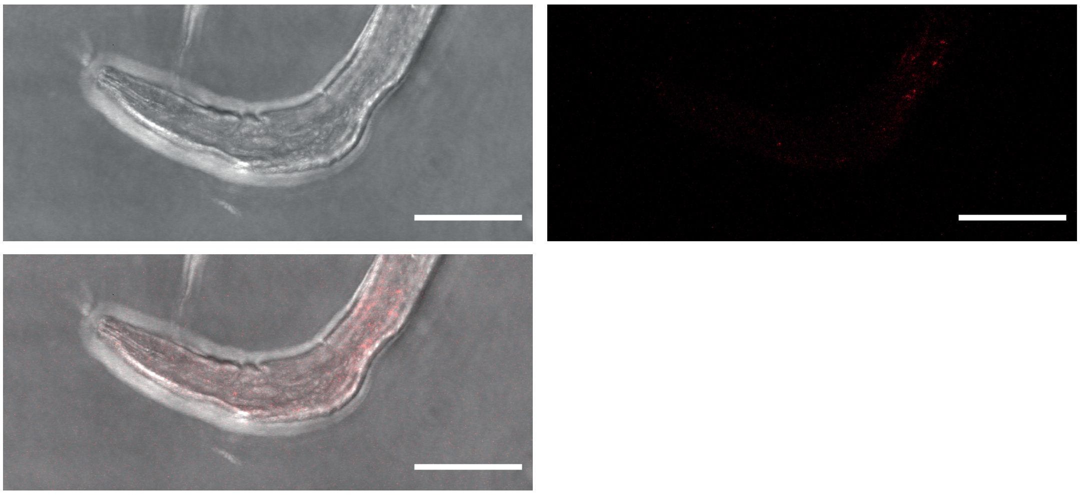

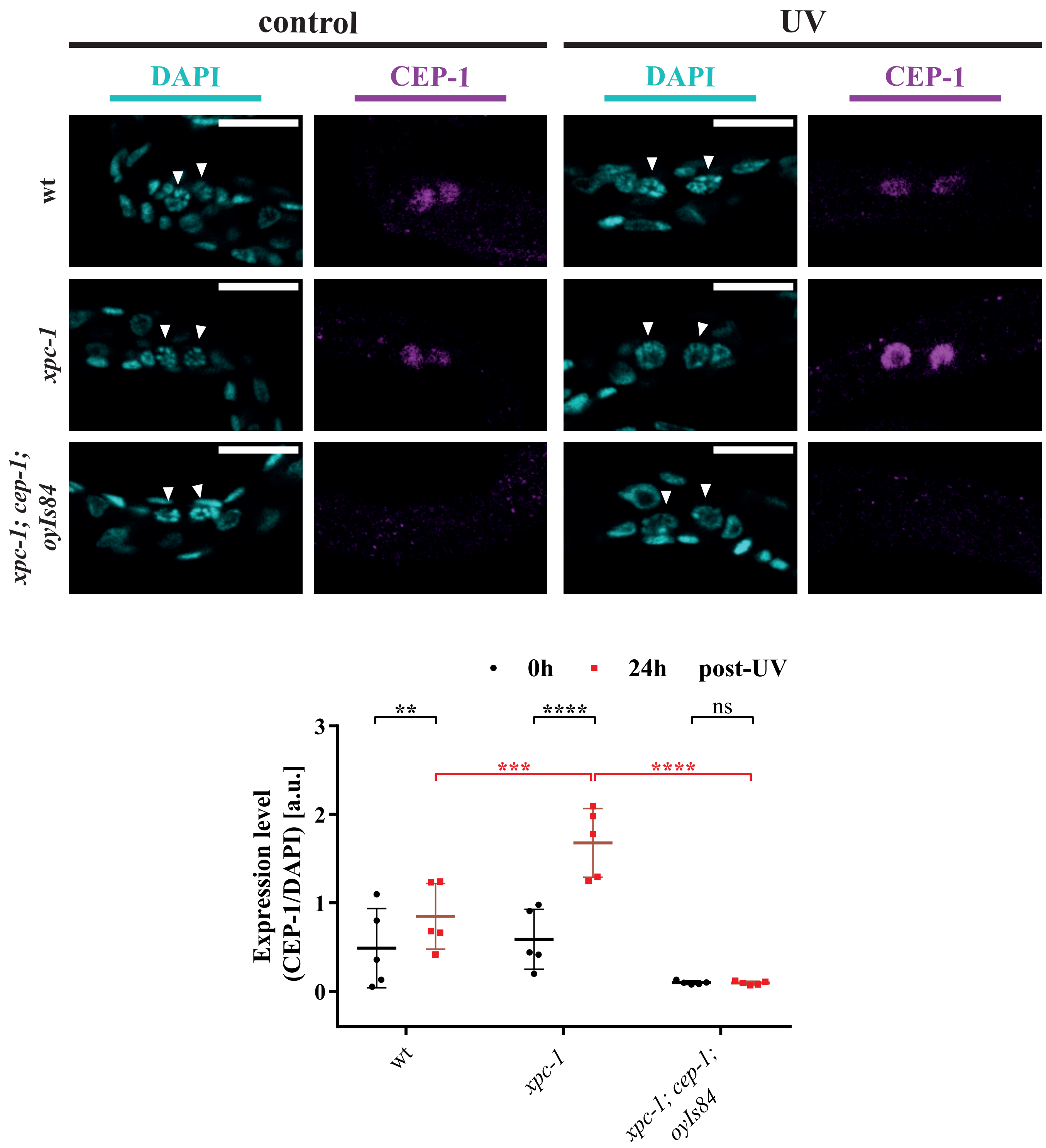
